# NLRP3-induced systemic inflammation controls the development of JAK2V617F mutant myeloproliferative neoplasms

**DOI:** 10.1101/2024.03.09.583936

**Authors:** Ruth-Miriam Koerber, Calvin Krollmann, Kevin Cieslak, Elisabeth Tregel, Tim H. Brümmendorf, Steffen Koschmieder, Martin Griesshammer, Ines Gütgemann, Peter Brossart, Radek C. Skoda, Carl Christian Kolbe, Eicke Latz, Dominik Wolf, Lino L. Teichmann

**Author notes:** corresponding author #**Correspondence** Lino L. Teichmann Department of Medicine III University Hospital Bonn Venusberg-Campus 1 Bonn, Germany, email address, Dominik Wolf Internal Medicine V, Comprehensive Cancer Center Innsbruck (CCCI) Medical University of Innsbruck, Anichstraße 35 Innsbruck, Austria. contributed equally (Dominik Wolf, Lino L. Teichmann).

## Abstract

The development of Philadelphia chromosome-negative classical myeloproliferative neoplasms (MPN) involves an inflammatory process that facilitates outgrowth of the malignant clone and correlates with clinical outcome measures. This raises the question to which extent inflammatory circuits in MPN depend on activation of innate immune sensors. Here, we investigated whether NLRP3, which precipitates inflammasome assembly upon detection of cellular stress, drives murine JAK2V617F mutant MPN. Deletion of *Nlrp3* within the hematopoietic compartment completely prevented increased IL-1β and IL-18 release in MPN. NLRP3 in JAK2V617F hematopoietic cells, but not in JAK2 wild type radioresistant cells, promoted excessive platelet production via stimulation of the direct thrombopoiesis differentiation pathway, as well as granulocytosis. It also promoted expansion of the hematopoietic stem and progenitor cell compartment despite inducing pyroptosis at the same time. Importantly, NLRP3 inflammasome activation enhanced bone marrow fibrosis and splenomegaly. Pharmacological blockade of NLRP3 in fully established disease led to regression of thrombocytosis and splenomegaly. These findings suggest that NLRP3 is critical for MPN development and its inhibition represents a new therapeutic intervention for MPN patients.

**Key points:**

- The increased IL-1β and IL-18 release in JAK2V617F mutant MPN depends on NLRP3 inflammasome activation
- NLRP3 in MPN promotes excess platelet production, granulocytosis, HSPC compartment expansion, splenomegaly and bone marrow fibrosis

## Introduction

The Philadelphia chromosome-negative, classical myeloproliferative neoplasms (for simplicity referred to as MPNs) are caused by somatic mutations in hematopoietic stem cells that promote clonal expansion of myeloid progenitor cells. MPNs encompass several clinicopathologic entities, i.e. polycythemia vera (PV), essential thrombocythemia (ET), and primary myelofibrosis (PMF). The diseases are characterized by myeloid cell proliferation, high risk of thromboembolism, constitutional symptoms and have the potential to progress to acute myeloid leukemia.

The gain-of-function V617F mutation in Janus tyrosine kinase 2 (JAK2) is the most common inducer of MPNs and is detected in 95% of PV and 50-60% of ET and PMF patients^1^. Notably, JAK2V617F has also been found in 3% of healthy individuals in a Danish general population study^2^. JAK2V617F causes constitutive STAT as well as PI3K/Akt and Ras/Raf/MAPK/ERK signaling. The oncogenic property of JAK2V617F resides in its ability to initiate ligand-independent signaling downstream of the erythropoietin receptor (EPOR), thrombopoietin receptor (TPOR), and granulocyte-stimulating factor receptor (G-CSFR), resulting in erythrocytosis, thrombocytosis, and neutrophilia, respectively^3^. In addition, JAK2 mediates signals from a multitude of other surface receptors, including receptors for chemokines, interleukins, interferons and receptor tyrosine kinases, many of which are critically involved in inflammatory responses^4^.

It is now well recognized that chronic inflammation impacts clinical outcome in MPN. TNFα has been shown to promote clonal dominance of JAK2V617F expressing cells by inhibiting healthy hematopoiesis^5^. Overexpression of IL-8 has been linked to constitutional symptoms, and that of HGF, CXCL9, and IL-1RA to splenomegaly^6^. PV and ET patients have higher serum concentrations of C-reactive protein than healthy individuals, which are associated with an increased incidence of cardiovascular events^7^. Most importantly, IL-8, IL-2R, IL-12, IL-15, and CXCL10 are independently associated with poorer overall survival in PMF^6^. Thus, inflammation in MPN is closely related to clonal selection, symptom burden, thromboembolism and survival.

Recent work has established IL-1β as a major regulator of inflammation in the aging hematopoietic stem cell niche^8^ and in MPN^9,10^. Deletion of *Il1b* in a JAK2V617F MPN mouse model reduced serum levels of a broad range of inflammatory cytokines^10^. Importantly, genetic or pharmacological inhibition of IL-1β or its receptor IL-1R1 was sufficient to ameliorate hallmark symptoms of MPN such as splenomegaly and myelofibrosis^9,10^. However, IL-1β is produced by cells as a cytosolic pro-form that depends on cleavage by the proinflammatory protease caspase-1 for its activity and extracellular release. Caspase-1 also matures IL-18 and gasdermin D, which can induce pyroptosis, a form of lytic cell death^11^. In addition it can degrade the transcription factor GATA1, which specifies erythroid cells and megakaryocytes^12^. Activation of caspase-1 requires the assembly of cytoplasmic multiprotein complexes termed inflammasomes^13^. The type of inflammasome driving MPN development remains to be identified.

Inflammasomes consists of a sensor, the adaptor molecule Apoptosis-associated speck-like protein containing a caspase recruitment domain (ASC) and the effector caspase-1. The best characterized inflammasome sensors are NLRP1, NLRP3, NLRC4, and AIM2. NLRP3 (NOD-, LRR-, and pyrin domain-containing 3) is essential for the immune response to numerous pathogens, but is also involved in chronic sterile inflammatory diseases, such as cryopyrin-associated periodic syndrome, type II diabetes, Alzheimer’s disease, atherosclerosis, and gout^14^. Recently, it has been implicated in the pathogenesis of myelodysplastic syndrome (MDS) ^15^ and KRAS mutant myeloid neoplasms^16^.

Here, we define the specific contributions of the inflammasome sensor NLRP3 to the pathogenesis of MPN by adopting two strategies, genetic deletion and pharmacological inhibition, to inactivate NLRP3 in a conditional JAK2V617F-driven mouse model of MPN.

## Results

### MPN patients exhibit increased spontaneous inflammasome activation

We performed cytokine profiling on serum samples from 173 MPN patients from the German Study Group MPN (GSG-MPN) bioregistry and 37 healthy controls. K-means clustering, an unsupervised algorithm, revealed a cluster of 17 cytokines (cluster A) that was upregulated in 43% of MPN patients but only in 13% of healthy controls (Fig. 1a). Notably, this cluster contained IL-1β and IL-18, whose maturation depends on caspase-1, suggesting inflammasome activation. The cluster also included IL-6, whose expression is induced by IL-1β, and TNFα, a priming factor for the NLRP3 inflammasome. We confirmed that IL-1β, IL-18 and TNFα levels were higher in MPN patients than in healthy controls when considered individually (Fig. 1b). In addition, serum concentrations of HMGB1, an alarmin that depends on inflammasome activation for its extracellular release, was also increased in MPN patients compared to healthy individuals (Fig. 1c). A hallmark of inflammasome activation is the ASC speck. We therefore quantified speck formation in peripheral blood mononuclear cells (PBMC) from MPN patients and healthy volunteers by flow cytometry^17^. The percentage of ASC speck positive cells in PBMCs without prior stimulation or after LPS plus nigericin was greater in MPN patients than in healthy individuals (Fig. 1d and 1e) and positively correlated with serum IL-1β concentrations (Fig. 1f). The increase in ASC speck positive cells was not due to a higher percentage of myeloid cells in MPN patients, as it was present in lymphocytes and myeloid cells (Fig. S1). Many of the PV and MF patients in this study were treated with the JAK inhibitor ruxolitinib, which exerts anti-inflammatory in addition to antimyeloproliferative effects. Notably, there was no difference in serum IL-1β and IL-18 concentrations between untreated PV and MF patients and those receiving ruxolitinib under real-world conditions (Fig. 1g).

**Fig. 1.**
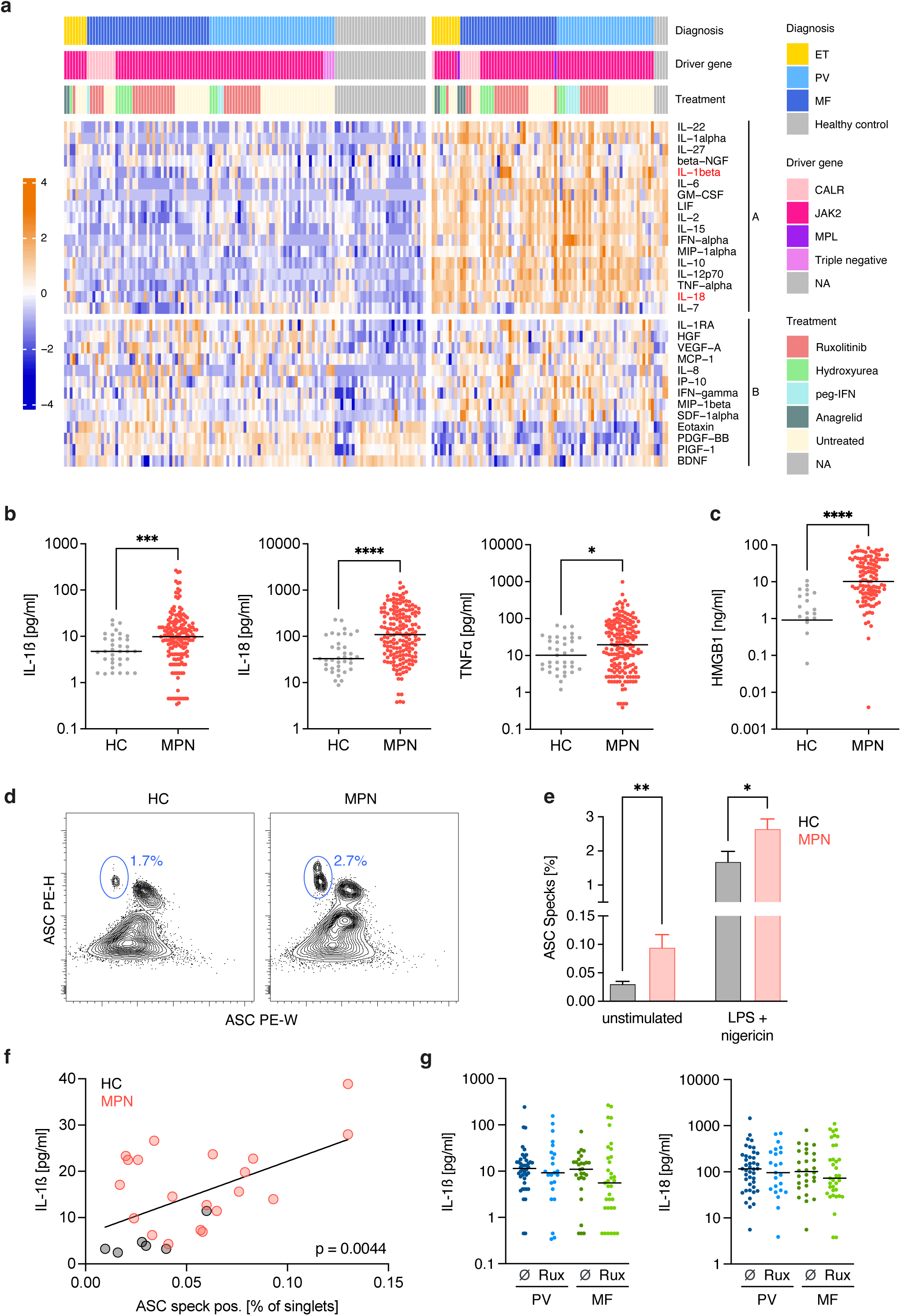
MPN patients exhibit increased spontaneous inflammasome activation. (a) Serum cytokines in healthy controls (n = 37) and MPN patients (n = 173) were measured by Luminex assay. Row- and column-wise K-means clustering with K = 2. Cytokine expression values are Z score standardized. Each column represents one individual. (b) IL-1β, IL-18 and TNFα serum concentrations in healthy controls (HC, n = 37) and MPN patients (n = 173). (c) HMGB1 serum concentration in healthy controls (HC, n = 30) and MPN patients (n = 129). (d) Representative ASC staining profiles and gating of ASC speck positive PBMCs in a healthy control (HC) and a MPN patient after LPS plus nigericin stimulation. Values indicate percentage of PBMCs (mean). (e) Percentage of ASC speck positive cells in PBMCs from healthy controls (HC, n = 14) or MPN patients (n = 36) without or after LPS plus nigericin stimulation. (f) Correlation of serum IL-1β concentration and percentage of ASC speck positive cells in PBMCs from healthy controls (HC, n = 6) and MPN patients (n = 20). (g) IL-1β and IL-18 serum concentration in ruxolitinib (Rux) treated and untreated (Ø) PV and MF patients. PV (Ø n = 42, Rux n = 23), MF (Ø n = 28, Rux n = 35) In scatter plots, each dot represents a single patient and horizontal lines the median. Bar graphs show mean + SEM. Statistically significant differences were determined by two-tailed unpaired Mann-Whitney U test (b, c, e, g) and Pearson correlation (f). \**P* < 0.05, \*\**P* < 0.01,\*\*\**P* < 0.001, \*\*\*\**P* < 0.0001.

Altogether, these findings demonstrate spontaneous inflammasome assembly in MPN patients.

### Caspase-1 dependent cytokine secretion in *Jak2^VF^* mice depends on the NLRP3 inflammasome

We next asked whether higher inflammasome activity in MPN patients is due to increased transcriptional priming. *IL1B* and *CASP1* mRNA was upregulated in PBMCs from MPN patients compared to those from healthy controls (Fig. 2a). To determine whether JAK2V617F promotes transcriptional priming in a cell-intrinsic fashion, we used a published scRNA-seq dataset of hematopoietic stem and progenitor cells (HSPCs) from MPN patients containing single cell genotype information^18^. JAK2V617F mutant HSPC showed stronger expression of an inflammasome gene signature than JAK2 wild type (WT) HSPCs from the same patients (Fig. 2b). These data support the concept that JAK2V617F cell-intrinsically upregulates inflammasome pathway components.

**Fig. 2.**
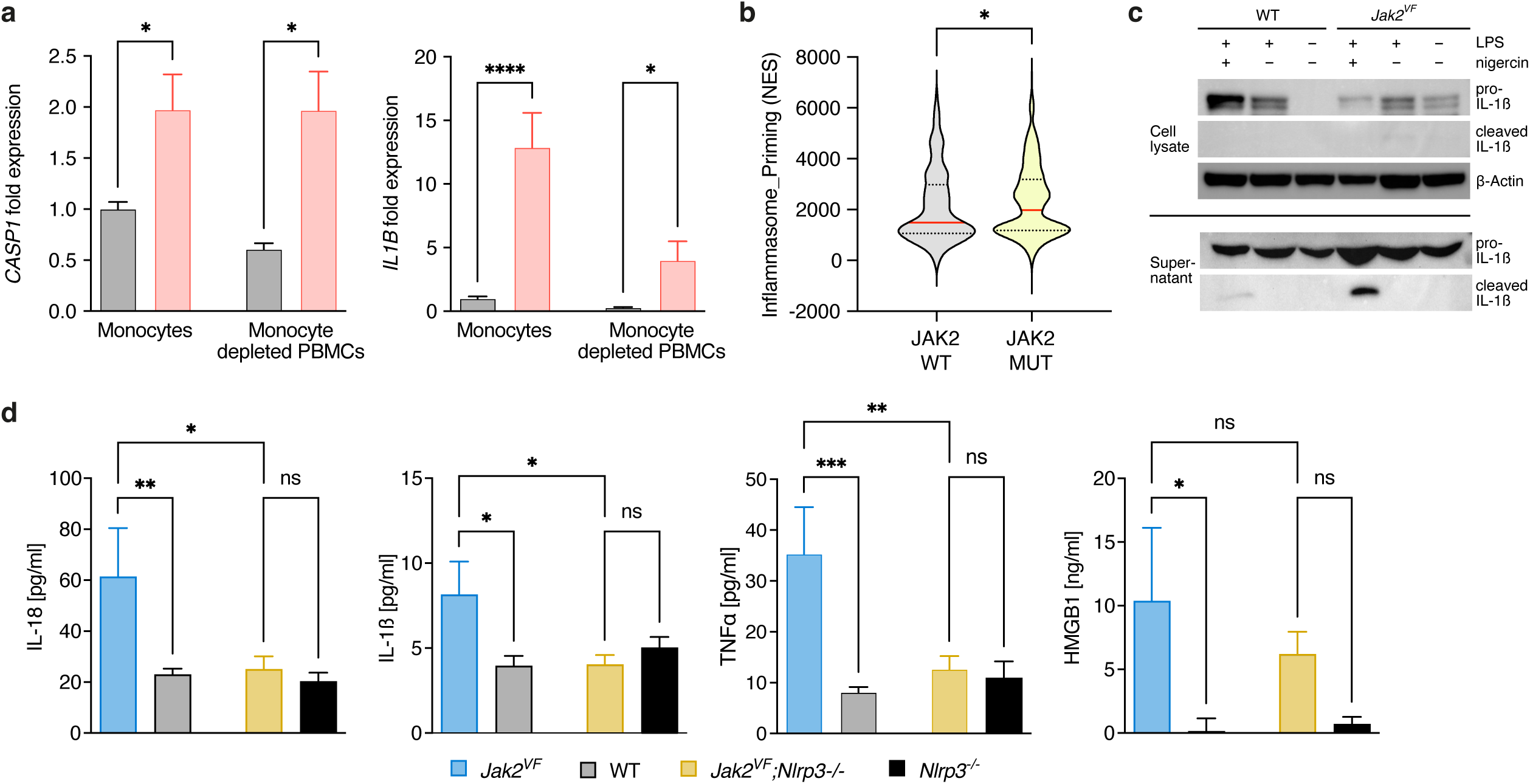
Caspase-1 dependent cytokine secretion in *Jak2^VF^* mice depends on the NLRP3 inflammasome. (a) *CASP1* and *IL1B* transcripts were quantified in monocytes and monocyte-depleted PBMCs from MPN patients (n = 52) and healthy controls (n = 14) by qRT-PCR. Data represent normalized expression values relative to monocytes from healthy controls. (b) Enrichment scores for a gene set comprising *IL1B, IL18, GSDMD, PYCARD* and *CASP1* (Inflammasome_Priming) in JAK2 WT and JAK2V617F mutant single HSPCs isolated from MF patients (GSE122198). (c) BMDMs from *Jak2^VF^*and WT mice were stimulated or left untreated as indicated, and cell lysates and supernatants were assessed for pro-IL-1ß and cleaved IL-1ß by immunoblotting. The blot is representative of 2 independent experiments. (d) IL-1β, IL-18, TNFα and HMGB1 serum concentrations in *Jak2^VF^*, WT, *Jak2^VF^;Nlrp3^-/-^* and *Nlrp3^-/-^* mice (each group n = 9) Bar graphs show mean + SEM. Violin plot displays median and quartiles. Statistically significant differences were determined by two-tailed unpaired Mann-Whitney U test (a and b) and one-way ANOVA with Holm-Šidák multiple comparison test (d). \**P* < 0.05, \*\**P* < 0.01,\*\*\**P* < 0.001, \*\*\*\**P* < 0.0001.

In the *Vav-Cre;Jak2^V617F/+^* mouse model (*Jak2^VF^*hereafter) of MPN, expression of JAK2V617F induces a PV-like phenotype^19^. We investigated whether inflammasome activation, as observed in MPN patients, also occurs in *Jak2^VF^* mice. Western blot analysis revealed higher amounts of pro-IL-1β in *Jak2^VF^*than WT bone marrow-derived macrophages (BMDM) even without any stimulation and cleaved IL-1β was reliably detectable in the supernatant of LPS plus nigericin stimulated BMDM from *Jak2^VF^* mice but barely evident in those from WT mice (Fig. 2c). IL-1β, IL-18, TNFα and HMGB1 serum levels were markedly elevated in *Jak2^VF^*relative to WT mice (Fig. 2d). To specifically test the role of the NLRP3 inflammasome in MPN we generated global NLRP3-deficient *Jak2^VF^* mice (*Jak2^VF^*;*Nlrp3^-/-^*). Strikingly, loss of NLRP3 in *Jak2^VF^* mice normalized IL-1β, IL-18 and TNFα serum concentrations (Fig. 2d).

Thus, in a JAK2V617F mutant mouse model of MPN, secretion of caspase-1 dependent cytokines relied on the NLRP3 inflammasome.

### NLRP3 drives thrombocytosis and granulocytosis in murine MPN

*Jak2^VF^* mice develop elevated erythrocyte, leukocyte and platelet counts^19^. We sought to investigate the impact of the NLRP3 inflammasome on blood cell production. *Jak2^VF^*, *Jak2^VF^;Nlrp3^-/-^* or WT bone marrow was transplanted into lethally irradiated WT mice (called *Jak2^VF^* BM, *Jak2^VF^;Nlrp3^-/-^*BM and WT BM mice hereafter) and blood was obtained by the submandibular vein method repeatedly over 25 weeks. Platelet numbers were greatly reduced in *Jak2^VF^;Nlrp3^-/-^* BM compared to *Jak2^VF^*BM mice; in fact, they were indistinguishable from those in WT BM mice (Fig. 3a). Hemoglobin and, for the most part, leukocyte levels were not affected by the deletion of *Nlrp3*. However, leukocyte counts are known to depend on the method of blood acquisition, with cardiac and vena cava bleeds providing the most consistent results^20^. Therefore, after 26 weeks, mice were sacrificed and blood from the abdominal vena cava was collected. In this analysis, platelets and leukocytes were both lower in *Jak2^VF^;Nlrp3^-/-^*BM than in *Jak2^VF^* BM mice (Fig. 3b). The drop in leukocytes was mainly due to a decrease in neutrophils, basophils, and eosinophils. Platelet leukocyte aggregates have been associated with thrombosis in MPN and are correlated with platelet counts^21^. As expected, neutrophil-platelet and inflammatory monocyte-platelet aggregates were decreased in *Jak2^VF^;Nlrp3^-/-^* BM relative to *Jak2^VF^*BM mice (Fig. 3c).

**Fig. 3.**
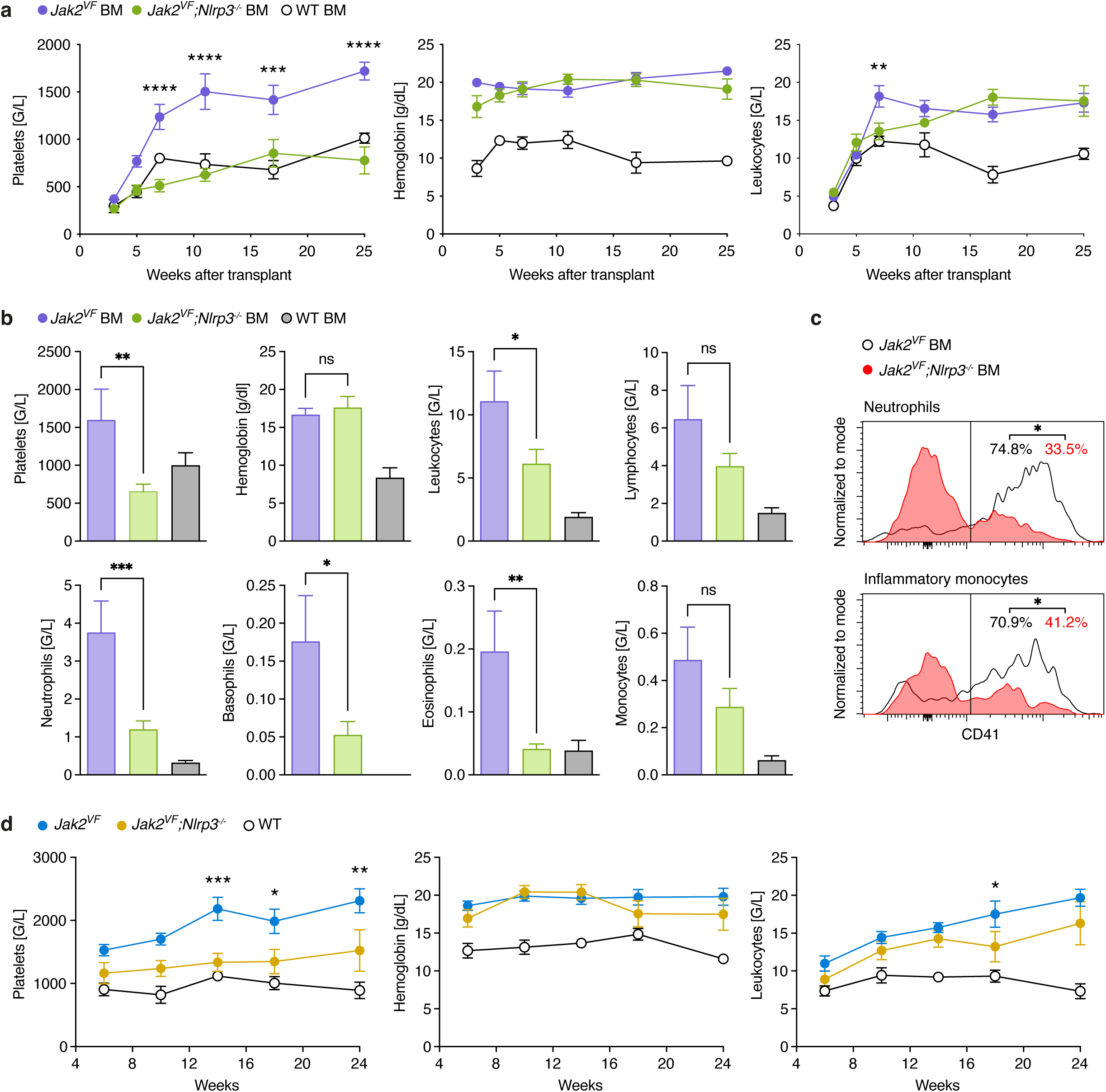
NLRP3 drives thrombocytosis and granulocytosis in murine MPN. (a) Blood counts of *Jak2^VF^* BM (n = 12), *Jak2^VF^;Nlrp3^-/-^* BM (n= 10) and WT BM (n = 6) mice. Blood was drawn by submandibular method. For clarity, only the significant differences between *Jak2^VF^*BM and *Jak2^VF^;Nlrp3^-/-^* BM are shown. (b) Differential blood count of *Jak2^VF^* BM (n = 5), *Jak2^VF^;Nlrp3^-/-^*BM (n = 9) and WT BM (n = 6) mice 26 weeks after bone marrow transplantation. Blood was collected from the vena cava. (c) Staining histograms display gating of leukocyte-platelet aggregates in *Jak2^VF^* BM (n = 5) and *Jak2^VF^;Nlrp3^-/-^* BM (n = 4) mice. Values indicate the mean percentage of neutrophil-platelet and inflammatory monocyte-platelet aggregates. (d) Blood counts of non-transplanted *Jak2^VF^* (n = 8), *Jak2^VF^;Nlrp3^-/-^* (n = 10) and WT (n = 9) mice Blood was drawn by submandibular method. The significant differences between *Jak2^VF^* and *Jak2^VF^;Nlrp3^-/-^* are displayed. Plots show mean + SEM. Statistically significant differences were determined by Mixed-effects model with Dunnett’s multiple comparisons test (a and d), one-way ANOVA with Holm-Šidák multiple comparison test (b) and two-tailed unpaired Mann-Whitney U test (c). \**P* < 0.05, \*\**P* < 0.01,\*\*\**P* < 0.001, \*\*\*\**P* < 0.0001.

A recent report suggested that NLRP3 may promote hematopoietic stem cell engraftment after transplantation^22^. In this study, platelets and leukocytes were reduced in mice transplanted with *Nlrp3^-/-^*bone marrow compared to those receiving WT bone marrow until 3 weeks after transplantation, but the difference disappeared after 4 weeks. Thus, an engraftment defect due to NLRP3 deficiency was an unlikely explanation for the lower platelet and leukocyte counts in *Jak2^VF^;Nlrp3^-/-^*BM than in *Jak2^VF^* BM mice, which persisted at least until 26 weeks after transplantation. Nevertheless, to investigate this possibility, we analyzed unmanipulated *Jak2^VF^* and *Jak2^VF^;Nlrp3^-/-^*mice (i.e. non-transplanted). *Jak2^VF^;Nlrp3^-/-^* mice showed lower platelets and granulocytes than *Jak2^VF^* mice, consistent with the results obtained in transplanted animals (Fig. 3d and Fig. S2a).

To investigate whether NLRP3 in non-hematopoietic radioresistant cells (such as epithelial and stromal cells) affects hematopoiesis in MPN we transplanted lethally irradiated WT and *Nlrp3^-/-^* mice with *Jak2^VF^* bone marrow. Lack of NLRP3 in radioresistant cells caused a slight reduction in platelets and leukocytes at week 5 after transplantation but had no effect beyond the engraftment period (Fig. S2b).

We conclude that *Nlrp3* deletion in hematopoietic cells impedes thrombocytosis and granulocytosis in JAK2V617F-induced murine MPN.

### The NLRP3 inflammasome promotes HSPC expansion and myeloid differentiation

JAK2V617F leads to an expansion of the HSPC compartment in MPN. The size of the LS-K cell population in the bone marrow containing CMPs (common myeloid progenitor), GMPs (granulocyte-monocyte progenitor), MEPs (megakaryocytic-erythroid progenitor), and MkPs (megakaryocyte progenitor) was diminished in *Jak2^VF^;Nlrp3^-/-^* BM relative to *Jak2^VF^* BM mice and comparable to that in WT BM mice (Fig. 4a and Fig. S3). Likewise, there were fewer LSK cells in the bone marrow of *Jak2^VF^;Nlrp3^-/-^* BM than of *Jak2^VF^* BM mice. All LSK subsets, i.e. HSCs, MPPs (multipotent progenitor) and HPCs (hematopoietic progenitor cells), in *Jak2^VF^;Nlrp3^-/-^*BM mice were reduced to the numbers observed in WT BM mice. We made largely similar findings in the spleen (Fig. 4b). Thus, the NLRP3 inflammasome drives enlargement of the HSPC pool in the bone marrow and spleen in murine MPN.

**Fig. 4.**
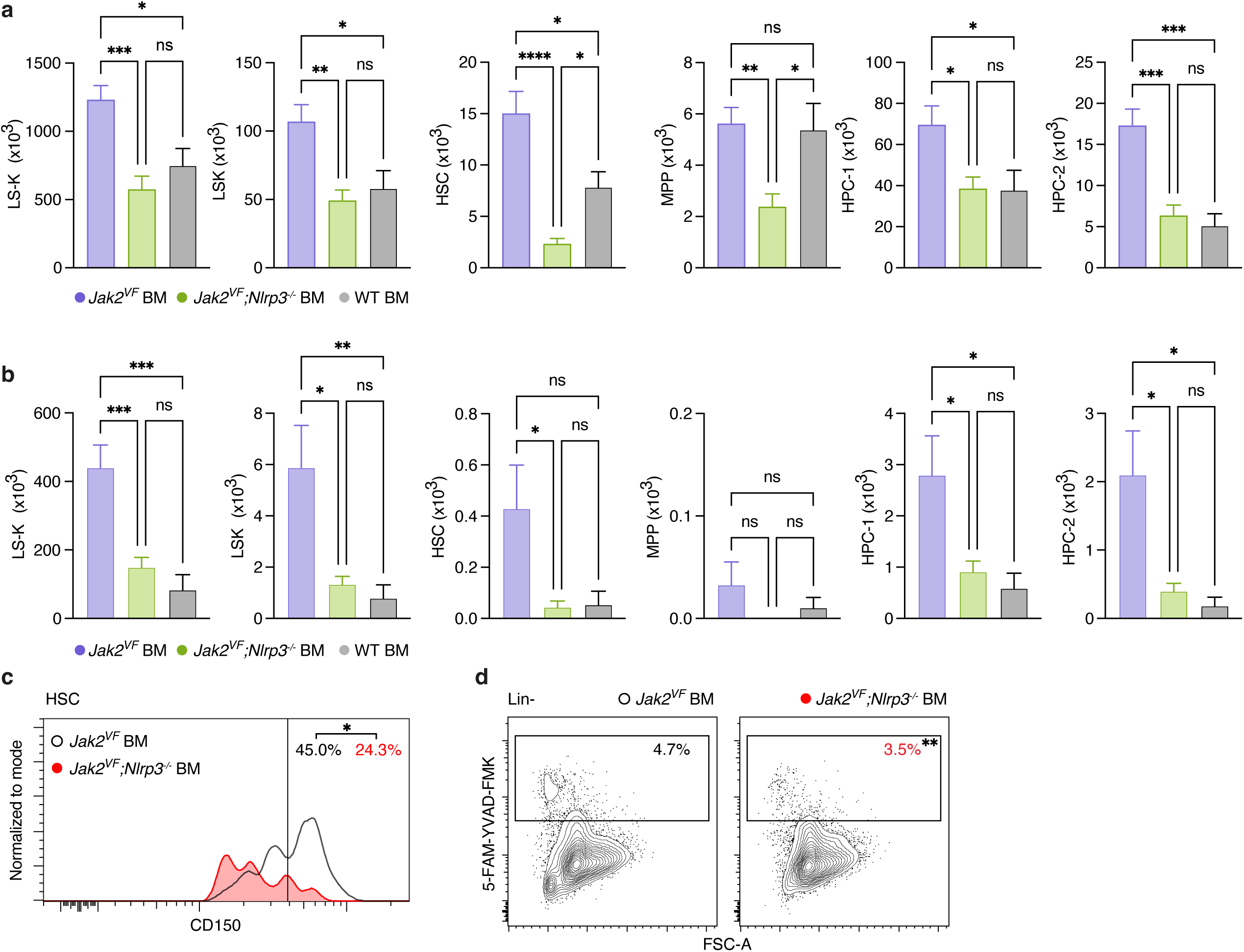
The NLRP3 inflammasome promotes HSPC expansion and myeloid differentiation. (a) HSPC subset counts in the bone marrow from both femurs of *Jak2^VF^* BM (n = 8), *Jak2^VF^;Nlrp3^-/-^* BM (n = 8) and WT BM (n = 6) mice. (b) As in (A) but in spleens of *Jak2^VF^* BM (n = 9), *Jak2^VF^;Nlrp3^-/-^*BM (n = 10) and WT BM (n = 5) mice. (c) Representative CD150 staining histograms of bone marrow HSCs in *Jak2^VF^* BM (n = 8) and *Jak2^VF^;Nlrp3^-/-^* BM (n = 8) mice. Values indicate the mean percentage of HSCs with high CD150 expression, indicating a myeloid bias. (d) Contour plots display gating of pyroptotic bone marrow HSPCs (Lin−) of *Jak2^VF^*BM (n = 8) and *Jak2^VF^;Nlrp3^-/-^* BM (n = 8) mice. Values indicate the mean percentage of HSPCs that undergo pyroptosis. Bar graphs show mean + SEM. Statistically significant differences were determined by one-way ANOVA with Holm-Šidák multiple comparison test (a and b) and two-tailed unpaired Mann-Whitney U test (c and d). \**P* < 0.05, \*\**P* < 0.01,\*\*\**P* < 0.001, \*\*\*\**P* < 0.0001.

HSCs represent a functionally heterogenous population. In MPN, HSCs show myeloid-biased differentiation. The expression level of CD150 on HSCs can be used to distinguish myeloid-from lymphoid biased HSCs^23^. Notably, deficiency for NLRP3 decreased the frequency of HSCs expressing high levels of CD150 in murine MPN (Fig. 4c), suggesting that NLRP3 imparts a myeloid skew, in keeping with published data on IL-1β^24^.

Pyroptosis is an inflammatory form of cell death caused by capase-1 cleavage of gasdermin D following inflammasome activation. We asked whether pyroptosis is induced in HSPCs by NLRP3. To identify cells undergoing pyroptosis we used the fluorescently labeled caspase-1/-4/-5 inhibitor 5-FAM-YVAD-FMK and scatter characteristics. Surprisingly, NLRP3 in JAKV617F mutant MPN indeed stimulated pyroptosis in HSPCs (Fig. 4d) even though it increased the HSPC population size.

Overall, NLRP3 promoted expansion and myeloid skewing of the HSPC pool in murine JAK2V617F-driven MPN.

### NLRP3 stimulates the direct thrombopoiesis pathway

NLRP3 deficiency potently suppressed platelet production in JAK2V617F-induced MPN. To better understand how NLRP3 regulates thrombopoiesis, we quantified megakaryocytes and MkPs in the bone marrow and spleen. Megakaryocytes were manually counted on tissue sections, as they tend to be underrepresented in flow cytometric assays because of their large cell size and fragility. *Jak2^VF^;Nlrp3^-/-^ z*BM mice contained substantially fewer megakaryocytes and MkPs than *Jak2^VF^* BM mice in bone marrow and spleen, respectively (Fig. 5a-c). Importantly, thrombopoietin serum concentrations were similar in *Jak2^VF^*BM and *Jak2^VF^;Nlrp3^-/-^*BM mice (Fig. 5d). Thus, NLRP3 deficiency did not lower platelet, megakaryocyte and MkP counts by reducing thrombopoietin levels.

**Fig. 5.**
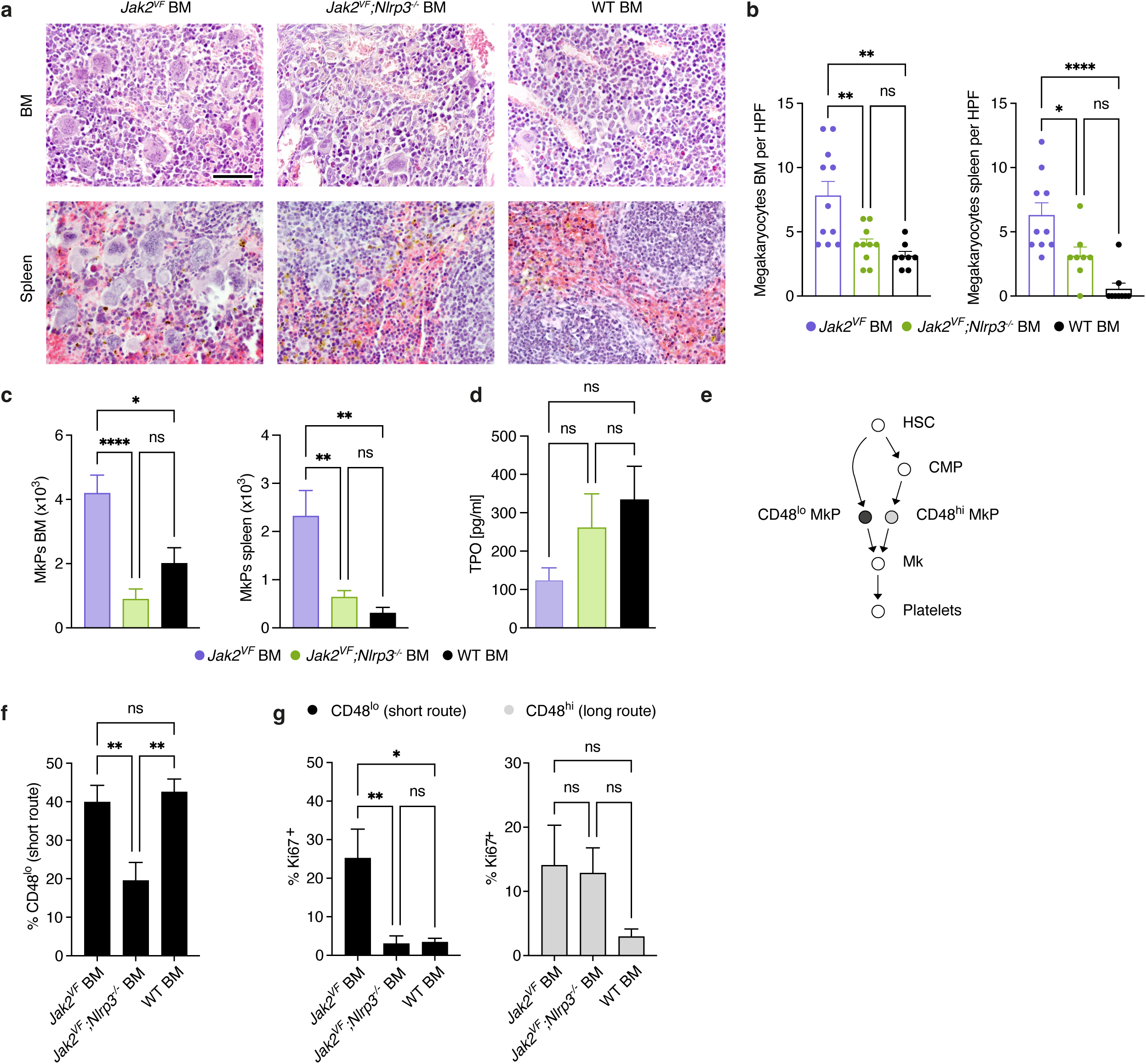
NLRP3 stimulates the direct thrombopoiesis pathway. (a) Representative images of H&E-stained bone marrow and spleen sections of *Jak2^VF^* BM, *Jak2^VF^;Nlrp3^-/-^* BM and WT BM mice at 26 weeks after bone marrow transplantation illustrating the megakaryocyte density. Scale bar equals 50 µm. (b) Megakaryocyte counts in *Jak2^VF^* BM, *Jak2^VF^;Nlrp3^-/-^* BM and WT BM mice (HPF, high-power field). Each dot represents a mouse. (c) MkP counts in *Jak2^VF^* BM (n = 8), *Jak2^VF^;Nlrp3^-/-^* BM (n = 8) and WT BM (n = 6) mice. (d) Thrombopoetin (TPO) serum concentration in *Jak2^VF^* BM (n = 11), *Jak2^VF^;Nlrp3^-/-^* BM (n = 10) and WT BM (n = 11) mice (e) Schematic of the direct and conventional thrombopoiesis pathway. (f) Percentage of MkPs that are CD48^lo^ in *Jak2^VF^* BM (n = 8), *Jak2^VF^;Nlrp3^-/-^*BM (n = 8) and WT BM (n = 6) mice. (g) Percentage of proliferating (Ki67^+^) CD48^lo^ and CD48^hi^ MkPs in *Jak2^VF^* BM (n = 8), *Jak2^VF^;Nlrp3^-/-^* BM (n = 8) and WT BM (n = 6) mice. Plots show mean + SEM. Statistically significant differences were determined by one-way ANOVA with Holm-Šidák multiple comparison test. \**P* < 0.05, \*\**P* < 0.01, **** *P* < 0.0001.

Until recently, the thrombopoiesis differentiation pathway was thought to proceed from HSCs via CMPs to MkPs. However, several studies have now provided evidence that megakaryocytes can also be replenished by direct differentiation of HSCs into MkPs^25–27^. Both, the long and the short route, are considered to be active simultaneously^26^. CD48 expression on MkPs distinguishes MkPs directly connected to HSCs (CD48^lo^ MkPs) from those that differentiate from CMPs (CD48^hi^ MkPs) (Fig. 5e). The frequency of MkPs that were CD48^lo^ was decreased in *Jak2^VF^;Nlrp3^-/-^* BM compared with *Jak2^VF^* BM mice (Fig. 5f), indicating that loss of NLRP3 impairs platelet production via the short route. Remarkably, deficiency for NLRP3 selectively reduced the frequency of actively cycling CD48^lo^ but not CD48^hi^ MkPs, as assessed by Ki67 staining (Fig. 5g).

These data demonstrate that NLRP3 stimulates the direct thrombopoiesis pathway in murine MPN, at least in part by inducing proliferation of associated CD48^lo^ MkPs.

### NLRP3 inflammasome activation propagates bone marrow fibrosis and splenomegaly

*Jak2^VF^* mice develop bone marrow fibrosis and, as a consequence of extramedullary hematopoiesis, splenomegaly. It remained unclear how NLRP3 would affect these major clinical manifestations of MPN. Because bone marrow transplantation delays the onset of fibrosis we assessed non-transplanted *Jak2^VF^*, *Jak2^VF^;Nlrp3^-/-^*and WT mice. Scoring of silver-stained tissue sections revealed that fibrosis development in the bone marrow and spleens of *Jak2^VF^* mice was greatly ameliorated in the absence of Nlrp3 (Fig. 6a and 6b). Accordingly, splenomegaly was also reduced in *Jak2^VF^;Nlrp3^-/-^* mice compared with *Jak2^VF^* mice (Fig. 6c). The attenuation of fibrosis caused by the lack of Nlrp3 was consistent with the lower number of megakaryocytes in *Jak2^VF^;Nlrp3^-/-^*mice, which are key drivers of bone marrow fibrosis^28^.

**Fig. 6.**
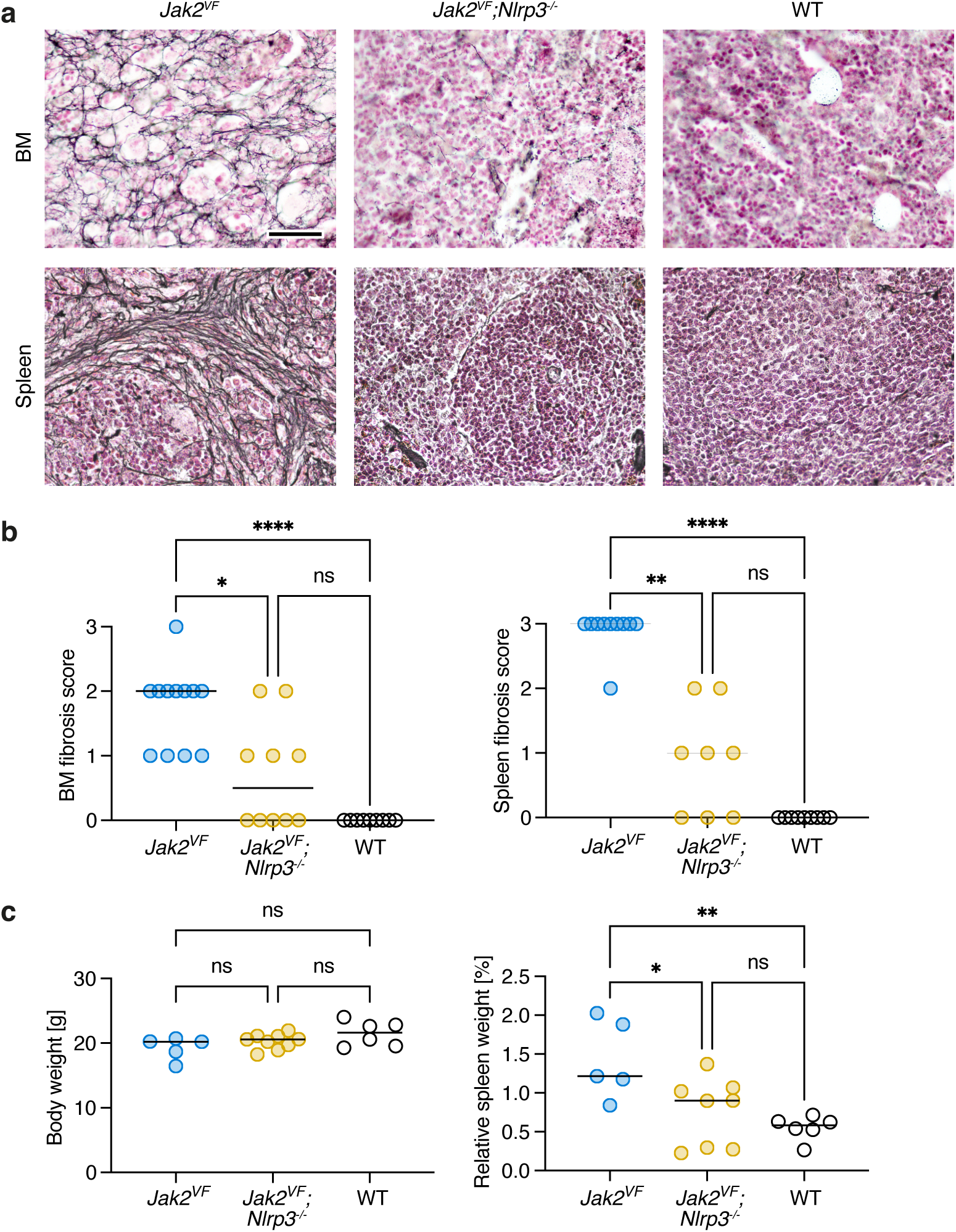
NLRP3 inflammasome activation propagates bone marrow fibrosis and splenomegaly. (a) Representative images of silver -stained bone marrow (femur) and spleen sections of *Jak2^VF^*, *Jak2^VF^;Nlrp3^-/-^*and WT mice at 26 weeks of age for grading of fibrosis. Scale bar equals 50 µm. (b) Fibrosis of bone marrow (left) and spleen (right) was scored from 0 to 3. Shown are scores of *Jak2^VF^*, *Jak2^VF^;Nlrp3^-/-^* and WT mice. (c) Body weight and relative spleen weight (percent of body weight). of *Jak2^VF^*, *Jak2^VF^;Nlrp3^-/-^* and WT mice In scatter plots each dot represents a mouse and bars the median. Statistically significant differences were determined by Kruskall-Wallis test with Dunn’s multiple comparison test (b) and one-way ANOVA with Holm-Šidák multiple comparison test (c). \**P* < 0.05, \*\**P* < 0.01, **** *P* < 0.0001.

### Pharmacological blockade of NLRP3 alleviates thrombocytosis, bone marrow fibrosis and splenomegaly

In the *Nlrp3* gene deletion studies, *Jak2^VF^* hematopoietic cells were already NLRP3-deficient in utero. We asked whether blocking NLRP3 after disease is already established would also lessen myeloproliferation. For this purpose, we used the novel oral NLRP3 inhibitor IFM-2384. *Jak2^VF^* BM mice received a chow formulation of IFM-2384 or control chow for 20 weeks starting 16 weeks after transplantation. On target efficacy was proven by reduced serum levels of IL-1β, TNFα and HMGB-1 in NLRP3 inhibitor-exposed *Jak2^VF^* BM mice (Fig. 7a). When blood was drawn from the submandibular vein, we found that in the IFM-2384 group, platelets decreased continuously from 914 G/L before to 333 G/L after 16 weeks of therapy (Fig. 7b). In contrast, thrombocytosis remained stable in the control group. There was also a trend for lower leukocytes in mice fed with IFM-2384 compared to those with control chow but this effect did not reach statistical significance.

**Fig. 7.**
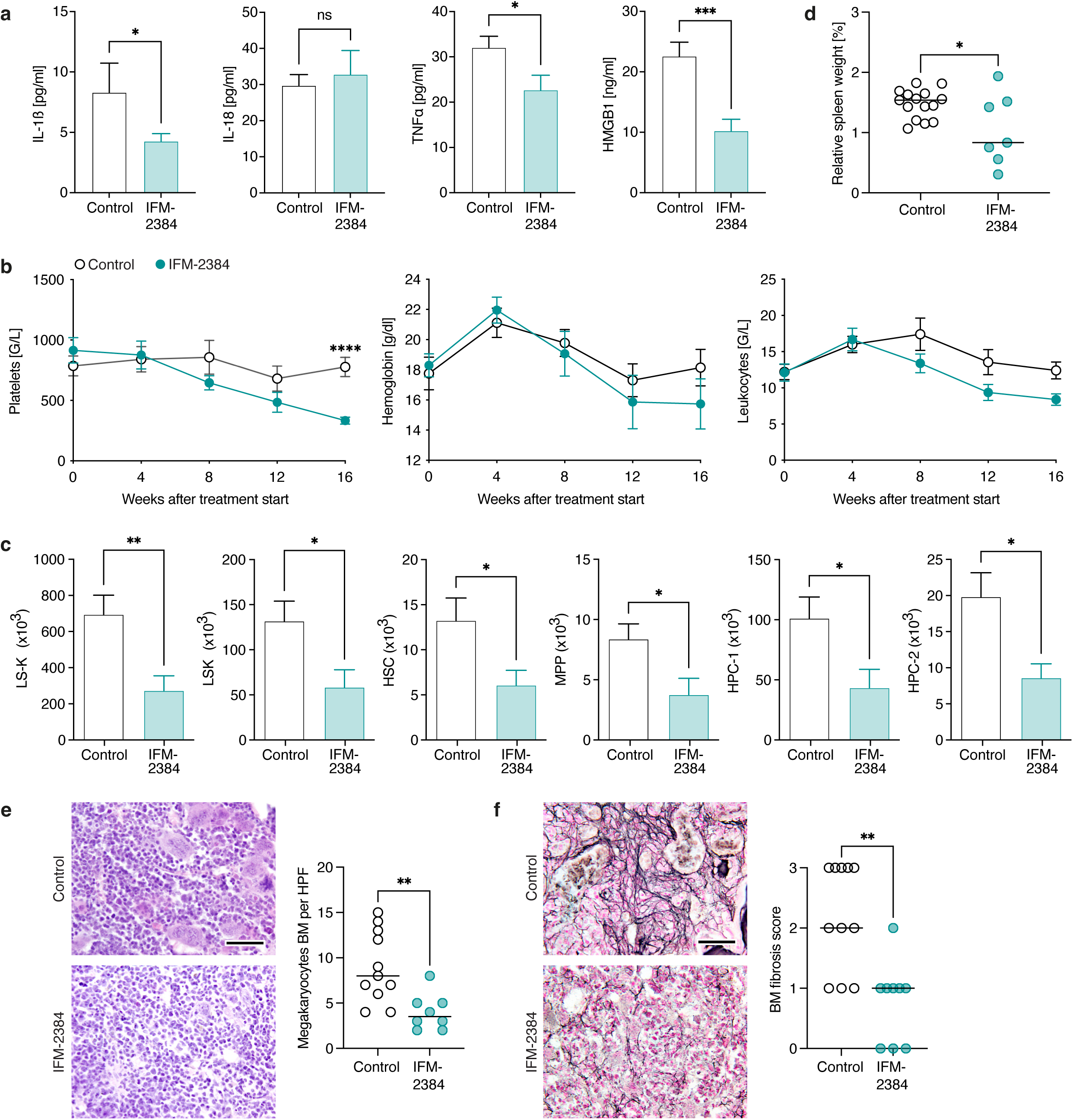
Pharmacological blockade of NLRP3 alleviates thrombocytosis and splenomegaly. *Jak2^VF^*BM mice were fed for 20 weeks with chow containing the oral NLRP3 inhibitor IFM-2384 or control chow starting at 16 weeks after transplantation. (a) IL-1β, IL-18, TNFα and HMGB1 serum concentrations in *Jak2^VF^* BM mice receiving IFM-2384 (n = 9) or control chow (n = 15). (b) Blood counts of *Jak2^VF^*BM mice fed IFM-2384 (n = 11) or control chow (n = 15). Blood was drawn by submandibular method. (c) HSPC subset counts in the bone marrow of *Jak2^VF^* BM mice receiving IFM-2384 (n = 11) or control chow (n = 15). (d) Relative spleen weight (percent body weight). Each dot represents a mouse. (e) Megakaryocyte counts in *Jak2^VF^* BM mice (right) with representative images of H&E-stained bone marrow sections (left). Each dot represents a mouse. Scale bar equals 50 µm. (f) Bone marrow fibrosis scores (right, scale 0 to 3) with representative images of silver-stained bone marrow sections (left). Each dot represents a mouse. Scale bar equals 50 µm. Graphs show mean + SEM except scatter plots that displays the median. Statistically significant differences were determined by two-tailed unpaired Mann-Whitney U test (a, c-f) and with unpaired t test with two-stage linear step-up procedure of Benjamini, Krieger and Yekutieli (b). \**P* < 0.05, \*\*\**P* < 0.001, **** *P* < 0.0001.

After 20 weeks of treatment, the mice were sacrificed and blood was obtained from the abdominal vena cava, which showed that NLRP3 inhibition decreased platelets and leukocytes, including neutrophils (Fig. S4). In keeping with the platelet data, there were fewer bone marrow megakaryocytes in *Jak2^VF^* BM mice fed IFM-2384 containing chow compared to those given control chow (Fig. 7e). Furthermore, IFM-2384 administration restricted HSPC expansion in murine MPN (Fig. 7c). Finally, NLRP3 blockade considerably mitigated splenomegaly and bone marrow fibrosis in *Jak2^VF^*BM mice (Fig. 7d and 7f).

These inhibitor studies further support the notion that NLRP3 is a viable therapeutic target in MPN.

## Discussion

Our study provides detailed insights into the essential contributions of the NLRP3 inflammasome to MPN development. By targeting NLRP3 genetically or pharmacologically in a JAK2V617F mutant mouse model, we formally establish that NLRP3 potently promotes malignant thrombocytosis and, to a lesser extent, granulocytosis. Notably, deletion of *Nlrp3* specifically impaired platelet production via the direct thrombopoiesis pathway. The loss of NLRP3 protected *Jak2^VF^* mice from fibrosis in the bone marrow and spleen, highlighting the importance of inflammasome activation in fibrogenesis. Splenomegaly, another key manifestation of MPN, was also markedly diminished in the absence of NLRP3.

The model of thrombopoiesis has recently been considerably advanced based on the finding that MkPs can arise directly from HSCs without first proceeding through CMPs^25–27^. In a mouse model of mutant *Calr*-driven ET, it was recently shown that megakaryocyte differentiation mainly occurred via a short route that directly links HSCs with MkPs^29^. Furthermore, single-cell transcriptomic data in MPN patients revealed that a direct differentiation route from HSCs to MkPs is aberrantly expanded compared with healthy individuals^30^. These studies provide evidence that the direct thrombopoiesis pathway may be the predominant contributor to increased platelet output in MPN. The drivers that increase platelet production via this trajectory in MPN have not been explored. Our study suggests, based on the differential expression of CD48 on MkPs, that the NLRP3 inflammasome is a major stimulator of the direct thrombopoiesis pathway in MPN.

NLRP3 might promote MPN development through several mechanism, including maturation of IL-1β and IL-18, initiation of pyroptosis by cleaving the pore-forming cell death executor gasdermin D and degradation of the hematopoietic lineage specifying transcription factor GATA1. Recently, the contribution of IL-1β and IL-1R1 were evaluated in MPN^9,10^. In these studies, deficiency for IL-1β or IL-1R1 attenuated bone marrow fibrosis and splenomegaly. These defects in the IL-1 pathway also impaired granulocyte and platelet production. In our work, NLRP3 exerted similar outcomes in JAK2V617F positive MPN, indicating that the phenotype of *Jak2^VF^* mice lacking NLRP3 is at least partly due to the reduction in mature IL-1β.

Inflammasome activation triggers pyroptosis via cleavage of gasdermin D by caspase 1. In MDS, the NLRP3 inflammasome has been implicated in the pathogenesis of cytopenia because it induced pyroptosis in HSPCs isolated from MDS patients^15^. We show that in MPN, NLRP3 also provokes pyroptosis in HSPCs but this did not cause cytopenia. On the contrary, NLRP3 rather promoted myeloproliferation. Therefore, other effects of NLRP3 appear to be more dominant over the impairment of hematopoiesis by pyroptosis in MPN. In the case of thrombopoiesis, it is noteworthy that IL-1β upregulates the megakaryocytic transcription factors NF-E2 and GATA1, as well as c-Fos and c-Jun, which are components of the dimeric activating protein-1 (AP-1) transcription-factor complex, thereby directly stimulating megakaryopoiesis^31^.

The transcription factor GATA1, which enforces megakaryocytic and erythroid differentiation, has been identified as a target of caspase-1^12^. Caspase-1 inhibition upregulates GATA1 protein in mouse HSPCs thereby promoting erythrocytosis. Degradation of GATA1 downstream of NLRP3 inflammasome activation has been proposed as a mechanism leading to anemia in MDS^32^. However, in our study of MPN in mice, NLRP3 had no effect on blood hemoglobin content and thus on erythrocytosis, although it was clearly activated. Overall, this highlights that the function of NLRP3 in MPN cannot readily be inferred from other diseases and its role has to be tested in vivo for each individual disease.

In addition to NLRP3, other inflammasomes may regulate malignant hematopoiesis in MPN. Expression of AIM2 is induced by JAK2V617F^33^ and in a mouse model of *Jak2* mutant clonal hematopoiesis, *Aim2* deficiency reduced atherosclerosis in an IL-1β-dependent manner^34^. Furthermore, NLRP1a has been found to trigger pyroptosis in HSPCs during hematopoietic stress caused by chemotherapy or viral infection leading to prolonged cytopenia^35^. We here focused on NLRP3 because its expression in increased in MPN patients^36^. We show that serum concentrations of IL-1β and IL-18 in *Jak2^VF^* mice lacking NLRP3 are reduced to those in WT mice. This does not exclude the possibility that other inflammasomes are involved in the development of MPN, but it does indicate that NLRP3 is an important inflammasome sensor in this disease.

Formation of the NLRP3 inflammasome complex is widely considered to involve a two-signal mechanism, with signal 1 required for transcriptional upregulation of NLRP3 and other inflammasome components (also known as ‘priming’). Subsequently, signal 2 can trigger NLRP3 inflammasome activation leading to caspase-1 activation. Signal 1 is typically induced by cytokine receptors or Toll-like receptor ligands, but the stimuli for signal 2 are more diverse and a unifying mechanism has yet to be defined^14^. Notably, numerous studies have now established that two signals are not always needed for assembly of the NLRP3 inflammasome, especially in vivo^37,38^. Our work reveals that JAK2V617 causes inflammasome priming, likely in a cell-intrinsic fashion. This is consistent with the fact that JAK2V617 induces NFκB signaling^39^. Whether a dedicated signal 2 is essential for NLRP3 activation in MPN remains to be elucidated in future studies. It is worth pointing out that in other myeloid malignancies reactive oxygen species (ROS) have been linked to NLRP3 activation^15,16^, but ROS inhibitors appear to rather block priming than activation^40^.

Our work has also limitations. In our mouse models all hematopoietic cells were of the *Jak2^VF^* genotype. Thus, it was not possible to assess whether NLRP3 activation promotes clonal outgrowth of JAK2V617 mutant cells. We also did not investigate the effect of the NLRP3 inflammasome on WT hematopoiesis. Because a Western-type calorically rich diet induces Nlrp3- dependent inflammation, we anticipate that such an analysis would need to take into account dietary conditions^41^.

In conclusion, we have delineated unique immunopathological effects of the NLRP3 inflammasome in JAK2 mutant MPN. Although more work is needed to define its regulation, our data confirm NLRP3 as a therapeutic target in patients with MPN, particularly those in whom excessive platelet production is of primary concern, such as in ET and prefibrotic PMF.

## Methods

### Primary human samples

MPN blood samples were obtained from patients enrolled in the German Study Group Myeloproliferative Neoplasms (GSG-MPN) bioregistry, approved by the Institutional Review Boards of the participating centers (DRKS-ID: DRKS00006035). Healthy control blood samples were drawn at the University Hospital of Bonn and collection was approved by the local Ethics Committee (154/13). Human studies were conducted according to the guidelines of the Declaration of Helsinki, and all subjects provided written informed consent before blood sampling.

### Isolation and processing of PBMCs

Whole blood was collected into EDTA tubes, and PBMCs were isolated by density gradient centrifugation with lymphocyte separation medium (PromoCell). PBMCs were either immediately used or cryopreserved at −80°C in RPMI1640 complemented with 20% FCS and 10% DMSO. Monocytes were isolated from PBMCs using the Pan Monocyte Isolation Kit (Milteny).

### Mice

*Vav-Cre;Jak2^V617F/+^* mice (*Jak2^VF^*) were kindly provided by Jean-Luc Villeval (Hasan et al. Blood, 2013). *Jak2^VF^;Nlrp3^-/-^* mice were generated by interbreeding *Jak2^VF^* with *Nlrp3^-/-^* mice (Millennium Pharmaceuticals). C57BL/6J mice were purchased from Jackson Laboratories. Mice were maintained under special pathogen free (SPF) conditions. Experimental procedures were performed in accordance with the German Animal Welfare Act and approved by the State Agency for Nature, Environment and Consumer Protection, NRW.

### Bone marrow transplantation

To generate bone marrow transplanted mice 10–12-week-old recipients were irradiated with 2x4.5 Gy in a 4h interval and injected intravenously with 2,5 x 10^6^ bone marrow cells harvested from 6–8-week-old donors. In all groups, care was taken to balance the ratio of female to male donors. Female mice were used as recipients. Analysis was performed 8-40 weeks after transplantation.

### Preparation of single-cell suspensions

To prepare bone marrow cells, femurs were extracted and soft tissue removed with a scalpel and by gentle rolling over paper towels. Femurs were cut open on one side and placed into a 0.5 ml tube with a small hole at the tip, which was inserted into another 1.5 ml tube. Bone marrow cells were flushed from the bone by centrifugation. For splenocytes, spleens were removed, minced and pressed through a metal strainer. All cells were treated with Red Blood Cell Lysis buffer (Biolegend) and suspensions were filtered through 70 μm Nitex nylon mesh before further use.

### Sampling of mouse blood and treatment of mice

Blood was collected from the submandibular vein in live animals or from the abdominal vena cava after sacrifice. Blood counts were measured on an auto hematology analyzer (Mindray BC-5000 Vet) using EDTA-anticoagulated blood. In the pharmacological studies, mice were either fed a chow-based formulation of the NLRP3 inhibitor IFM-2384 (0,18 mg/g, IFM Therapeutics) or a control diet for 20 weeks.

### In vitro NLRP inflammasome activation

Cells were primed with LPS for 3 hours or overnight at 37 °C with 100-200 ng/ml LPS in RPMI1640 containing penicillin, streptomycin and 10% FCS. To activate NLRP3, nigericin (10 µM, Sigma Aldrich) was added to the cultures for 1h. For detection of ASC specks, caspase-1 was inhibited with belnacasan (50 µM, Selleckchem) prior to stimulation with nigericin.

### Multiplex bead-based assays and ELISA

Human cytokines were measured with a custom ProcartaPlex Immunoassay (Thermo Fisher Scientific) and murine cytokines by LEGENDplex assay (Biolegend). The ProcartaPlex assay was read on a Luminex FlexMAP 3D System and LEGENDplex assays on a FACS Canto II (BD). Human and murine HMGB-1 was quantified by ELISA (IBL).

### Quantitative RT–PCR

RNA was extracted from cells with the Maxwell RSC simplyRNA Cells Kit (Promega) on a Maxwell RSC instrument and mRNA reverse transcribed with the RevertAid First Strand cDNA Synthesis Kit (Thermo Fisher Scientific). *GAPDH* was used as housekeeping gene. qPCR was performed using the SensiFast Sybr No-Rox Mix (Bioline) on a RealPlex 2 Thermocycler (Eppendorf).

### Histology

Tissues were formalin-fixed, embedded in paraffin and sectioned (4-6 µm). Megakaryocytes were quantified in a blinded fashion on hematoxylin and eosin-stained sections. Fibrosis was scored in a blinded fashion on Gomori stained sections (0 = very little scattered linear reticulin with no intersections corresponding to normal BM, 1 = loose network of reticulin with few intersections, 2 = diffuse and dense increase in reticulin with extensive intersections, focal thick collagen fibers, 3 = coarse bundles of collagen fibers).

### Quantification and statistical analysis

Statistical analysis was performed with GraphPad 9 (Prism). The statistical tests used are indicated in the figure legends. Independent experiments were pooled and analyzed together whenever possible as detailed in figure legends. Clustering analysis and visualization was performed using the ComplexHeatmap package (2.14.0) and ssGSEA using the escape package (1.8.0) in R.

Antibody clones, primer sequences and further methods are provided in the Supplementary information file.

## Acknowledgments

We thank HET (Haus für Experimentelle Therapie) for outstanding animal husbandry, Alexander Gerbaulet for helpful discussion and critical reading of the manuscript, and the GSG-MPN centers for patient samples. This study was funded by the Deutsche Krebshilfe (DKH, German Cancer Aid) Mildred Scheel Early Career Center grant 70113307 (RMK) and the Deutsche Forschungsgemeinschaft (DFG, German Research Foundation) grant TRR237-369799452 (LLT). It was further supported by the DFG grant EXC2151–390873048 (LLT) under Germany’s Excellence Strategy, the Jose-Carreras-Leukämie Stiftung grant 16R/2018 (DW), and in part by DFG grants 417911533 (THB) and 327211770 (SK) within the clinical research unit CRU344.

## Authorship contributions

RMK and LLT designed experiments, analyzed the data and wrote the manuscript. RMK, CK, KC and ET performed experiments. CCK performed the Luminex assay. THB, SK and MG provided patient samples. IG helped with histology. PB and EL provided resources. RCS provided feedback on and edited the manuscript. DW and LLT conceived and supervised the study. All authors had the opportunity to discuss the results and comment on the manuscript.

