## Supplementary Information for "NLRP3-induced systemic inflammation controls the development of JAK2V617F mutant myeloproliferative neoplasms"

### Supplementary Methods

#### Antibody clones for flow cytometry

CD3 (clone 17A2, Biolegend), CD19 (clone 6D5, Biolegend), Ter-119 (clone Ter119, Biolegend), Gr-1 (clone RB6-8C5, Biolegend), CD34 (clone RAM34, invitrogen), CD117 (clone ACK2, Biolegend), Sca-1 (clone D7, invitrogen), CD41 (clone MWReg30, Biolegend), CD150 (clone TC15-12F12.2, Biolegend), CD48 (clone HM48-1, Biolegend), Ly6C (clone HK1.4, Biolegend), Ly6G (clone 1A8, Biolegend), CD11b (clone M1/70, Biolegend), CD11c (clone N418, Biolegend), Trustain (clone 93, Biolegend), EPCR (clone RCR-16, Biolegend), ASC (clone HASC-71, Biolegend), VCAM (clone M/K-2, Invitrogen), ICAM (clone YN1/1.7.4., Biolegend), Ki-67 (clone 16A8, Biolegend), CD42b (clone Xia.65, Emfret), GbVI (clone FAB6758G, R&D)

#### Flow cytometry

Surface staining was performed on ice in PBS containing 0.5% BSA and 0.05% sodium azide. Intracellular staining was carried out using the FoxP3 / Transcription Factor Staining Buffer Set (Thermo Fisher Scientific). For exclusion of dead cells ethidium monoazide bromide (EMA) or ZOMBIE dyes (Biolegend) were used. Activated caspase-1/4/5 was detected with the 5-FAM-YVAD-FMK kit (AAT Bioquest). Data were acquired on a FACS Canto II (BD) and analyzed with FlowJo 10.8.1 (BD).

#### Primer sequences (5’-3’)

*GAPDH* forward GAGTCAACGGATTTGGTCGT

*GAPDH* reverse TTGATTTTGGAGGGATCTCG

*IL1B* forward TGGGCAGACTCAAATTCCAGCT

*IL1B* reverse CTGTACCTGTCCTGCGTGTTGA

*CASP1* forward ACAACCCAGCTATGCCCACA

*CASP1* reverse GTGCGGCTTGACTTGTCCAT

**Generation of bone-marrow derived macrophages (BMDMs)**

$7x10^6$ bone marrow cells were cultured in tissue-culture-treated petri dishes in 8 ml of IMDM supplemented with glutamine, penicillin, streptomycin, 2-mercaptoethanol (all from Thermo Fisher Scientific), 10% FCS (Merck) and M-CSF (20 ng/ml, Biolegend). On day 4, 4 ml of fresh, prewarmed complete medium was added. On day 7, supernatants with non-adherent cells were discarded and adherent cells were carefully detached with a cell scraper and washed with PBS.

**Western Blot**

Whole cells lysates from BMDMs were prepared in RIPA buffer (Sigma-Aldrich) containing phosphatase inhibitor cocktails 2&3 (Sigma-Aldrich). Supernatant proteins were precipitated with chloroform/methanol. Protein concentrations were determined by bicinchoninic acid assay (Pierce). 40 µg of whole-cell lysates or precipitates of 500 µl supernatant were separated on a polyacrylamide gel and transferred on poly-vinyl dinitrofluoride (PVDF) membranes. PVDF membranes were then immunoprobed with antibodies against murine IL-1ß (DY401, R&D) and ß-Actin (#4967, Cell Signaling) followed by HRP-labeled secondary antibodies. Proteins were detected with an ECL system (GE Healthcare) on a ChemiDoc XRS+ (Bio-Rad).

### Supplementary Figures

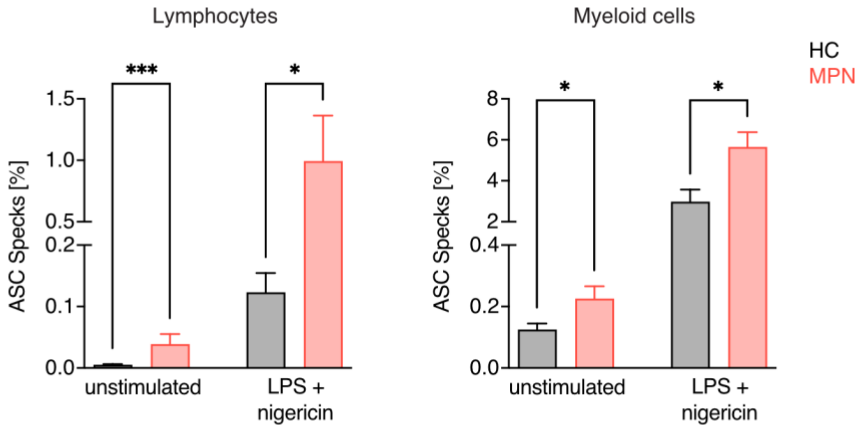


**Fig. S1. ASC speck formation is increased in MPN patients.**

Percentage of lymphocytes and myeloid cells with aggregated ASC in PBMCs from healthy controls (HC, n = 14) or MPN patients (n = 36) without or after LPS plus nigericin stimulation. Bar graphs show mean + SEM. Statistically significant differences were determined by two-tailed unpaired Mann-Whitney U test. *$P < 0.05$, ***$P < 0.001$.

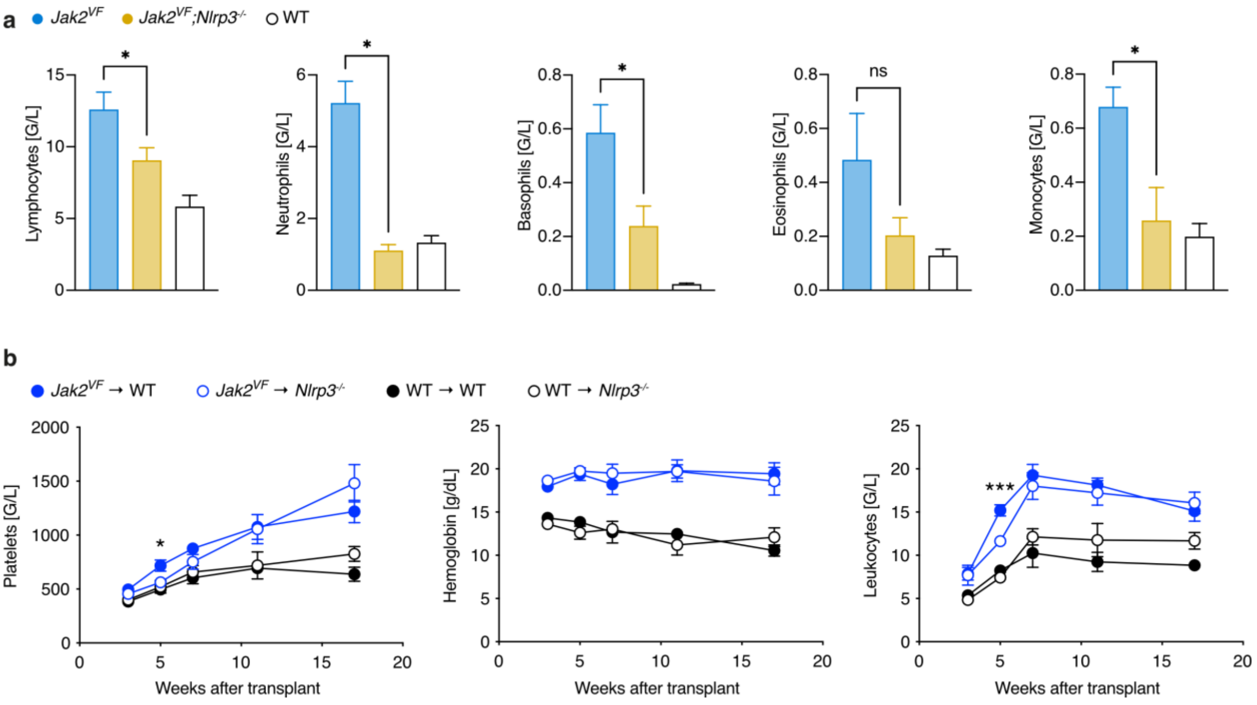


**Fig. S2. NLRP3 in radioresistant cells has no sustained effect on blood counts in murine MPN**

(a) Differential blood counts of non-transplanted *Jak2*$^{VF}$ (n = 8), *Jak2*$^{VF}$*;Nlrp3*$^{-/-}$ (n = 10) and WT (n = 9) mice at 25 weeks of age. Blood was collected from the abdominal vena cava. The significant differences between *Jak2*$^{VF}$ and *Jak2*$^{VF}$*;Nlrp3*$^{-/-}$ are displayed.

(b) Blood counts of lethally irradiated WT and *Nlrp3*$^{-/-}$ mice transplanted with *Jak2*$^{VF}$ or WT bone marrow (each group n = 10). Blood was drawn by submandibular method. For clarity, only the significant differences between *Jak2*$^{VF}$ → WT and *Jak2*$^{VF}$ → *Nlrp3*$^{-/-}$ mice are shown.

Plots show mean + SEM. Statistically significant differences were determined by one-way ANOVA with Holm-Šidák multiple comparison test (a) and mixed-effects model with Dunnett's multiple comparisons test (b). *$P$ < 0.05, ***$P$ < 0.001.

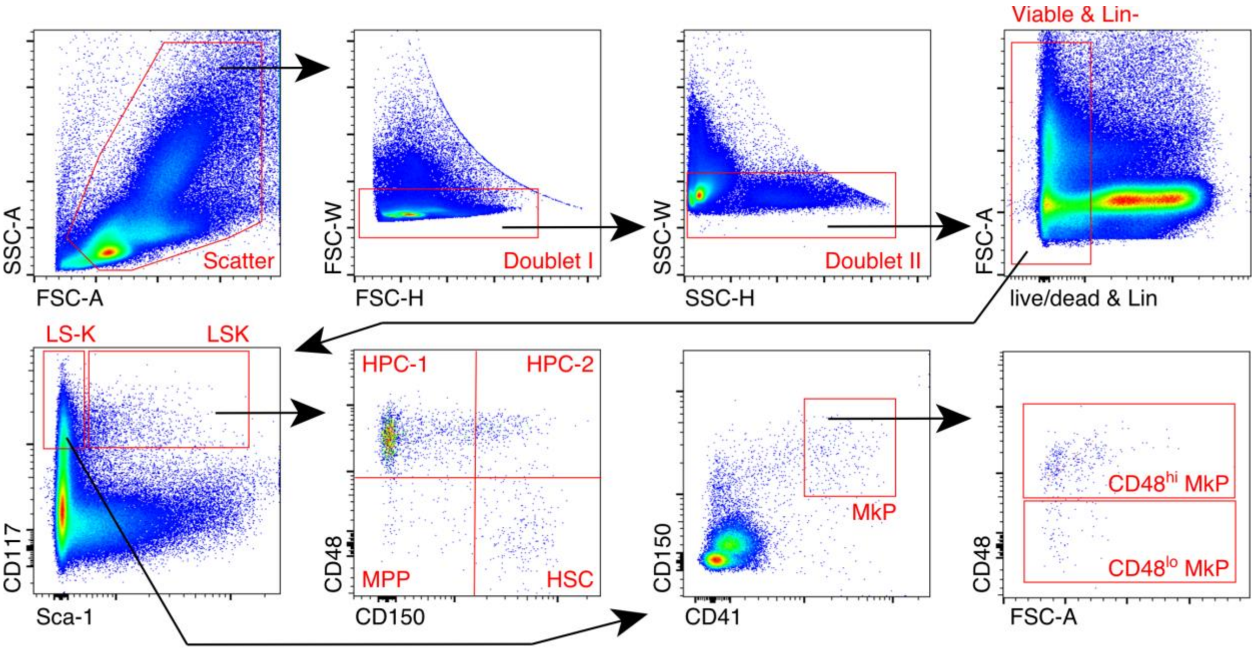


**Fig. S3. Flow cytometric identification of bone marrow HSPCs.**

Representative example of gating for bone marrow HSPCs. LS-K (Lin−, Sca-1−, CD117+), LSK (Lin−, Sca-1+, CD117+), HSC (Lin−, Sca-1+, CD117+, CD150+, CD48−), MPP (Lin−, Sca-1+, CD117+, CD150−, CD48−), HPC-1 (Lin−, Sca-1+, CD117+, CD150−, CD48+), HPC-2 (Lin−, Sca-1+, CD117+, CD150+, CD48+), MkP (Lin−, Sca-1−, CD117+, CD150+, CD41+).

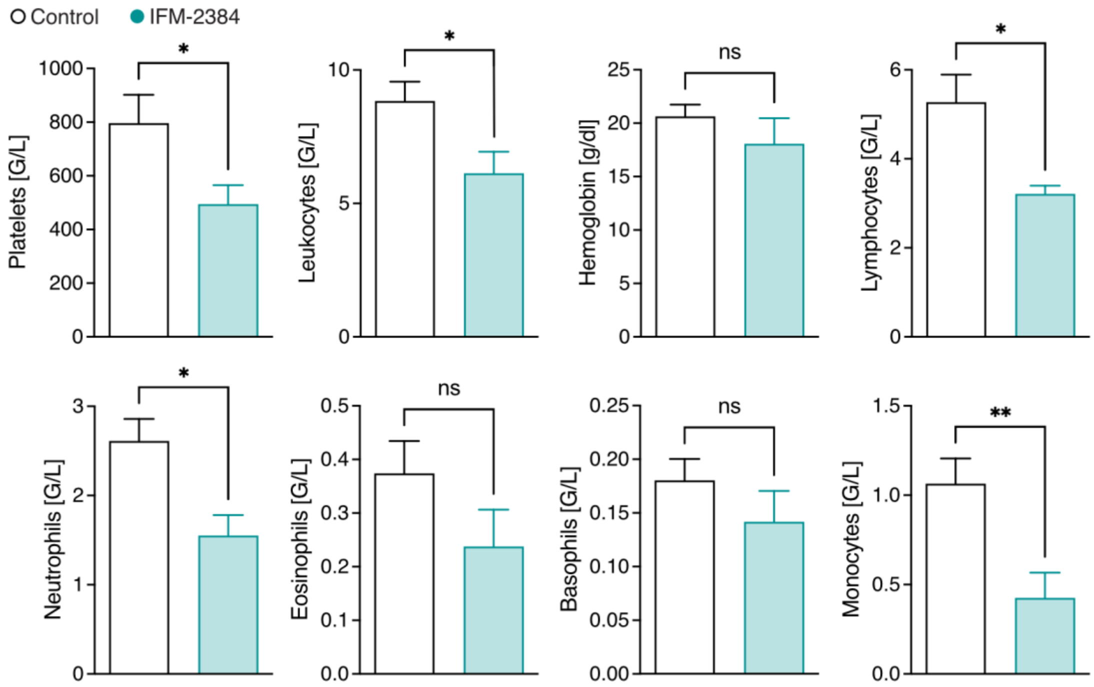


**Fig. S4. NLRP3 inhibition decreases platelets and leukocytes, including neutrophils, in *Jak2$^{VF}$* BM mice.**

Differential blood counts of *Jak2$^{VF}$* BM mice fed IFM-2384 (n = 11) or control chow (n = 15) 20 weeks after treatment start. Blood was collected from the abdominal vena cava.

Graphs show mean + SEM. Statistically significant differences were determined by two-tailed unpaired Mann-Whitney U test. *$P$ < 0.05, **$P$ < 0.01.
